# Characterising Type I Collagen Architecture in Human Bone and Osteosarcoma with Second Harmonic Generation Microscopy

**DOI:** 10.1101/2025.07.23.666332

**Authors:** Belle Creith, James Harrison, Peter B Johnson, Richard OC Oreffo, Claire E Clarkin, Sumeet Mahajan

**Author notes:** Equally contributing (Claire E Clarkin, Sumeet Mahajan).

## Abstract

Aberrant collagen matrix production is a hallmark of many cancers, including osteosarcoma (OS), the most common primary cancer of bone which remains poorly diagnosed. Here, the phenotyping potential of second harmonic generation (SHG) microscopy has been demonstrated through label-free imaging of pathologically altered collagen matrices within clinical biopsies. This work examined SHG imaging in quantifying OS-specific collagen (type I) signatures – focused on fibre length-based parameters – across clinical biopsies. We describe a novel SHG analysis workflow, following systematic optimisation of image resolution and field of view (FOV) size to enable robust quantification of collagen fibre-length metrics in human bone and OS. This included analyses of SHG images with a FOV spanning 350 µm × 350 µm to 1400 µm × 1400 µm, with the latter enabling whole-biopsy visualisation via stitching of up to 64 regions. In normal cortical (rib) bone biopsies, average fibre lengths did not significantly differ across increasing FOV size (26.44 ± 1.75 µm, 26.34 ± 2.01 µm, 27.25 ± 2.47 µm, 28.78 ± 3.58 µm). However, analysis of the top 50th percentile of collagen fibres revealed significant differences between the smallest FOVs; 350 µm × 350 µm and 466 µm × 466 µm, versus the largest 1400 µm × 1400 µm (p = 0.03 and p = 0.04 respectively), highlighting the requirement for larger FOVs for optimal characterisation of human bone collagen. Comparison of normal bone and OS (stage IIB) biopsies, the largest FOV revealed the largest differences, with OS exhibiting significantly reduced collagen fibre lengths (female OS: 23.18 ± 0.56 µm vs. bone: 29.39 ± 2.17 µm, p < 0.001; male OS: 24.39 ± 1.32 µm vs. bone: 27.87 ± 1.05 µm, p = 0.02) versus normal bone. Subsequent analyses across OS stages (IB – IVB) showed further evidence of pathological collagen signatures amplified with OS progression. These findings demonstrate that SHG microscopy enables robust, quantitative phenotyping of human bone and osteosarcoma collagen matrices, offering new potential for improved diagnosis and disease staging.

## Introduction

Clinical examination of the mineralised skeleton presents many challenges, including effective assessment of micro and nano-structural properties contributing to bone pathologies such as cancer progression and metastasis. These constraints exist, in part, due to limitations of current imaging modalities in resolving early, cellular level disease manifestations, which are typically concealed deep within the mineralised bone matrix. As a result, challenges remain in the diagnosis of osteosarcoma (OS), the most common primary cancer of bone, manifesting as aggressive paediatric tumours associated with aberrant production of non-mineralised extracellular matrix (ECM) by neoplastic cells of mesenchymal origin [1].

Tumorigenic remodelling of the ECM can aid metastasis, alter matrix rigidity and vessel permeability, influencing processes such as drug delivery and cyto-toxic therapy resistance – thus, contributing to cancer advancement [2]. The significant role of collagen in regulating key physiological processes such as cell proliferation, cell migration and immunity, is also exploited by many tumours, including OS, to promote vascularisation and tumour growth [3, 4]. Moreover, the architectural organisation of collagen has been shown to profoundly influence neoplastic progression, with fibrillar collagen inducing cellular behaviours characteristic of invasive tumours [5]. In healthy tissues, including bone, the ECM plays a crucial role in shaping structural integrity and function. Existing as the most abundant matrix protein, type I collagen, provides mechanical integrity to bone whilst influencing cellular behaviour and signalling [6].

Today, successful diagnoses and treatment advances for OS remains poor, highlighting an urgent need for innovative early diagnostic technologies to be developed. In recent years, non-linear microscopy has emerged as an alternative approach for the examination and potential diagnosis of cancers – predominantly in soft tissues – through detection of disease-specific ECM production, as reviewed in [7]. Non-linear microscopy refers to a suite of optical imaging techniques that utilise pulsed laser sources to provide complementary readouts of molecular structures within tissues, providing histology-like images [8]. Numerous non-linear microscopy techniques are intrinsically label-free, exploiting optical phenomena that generate contrast for imaging without need for exogenous labelling and laborious sample preparation. Second harmonic generation (SHG) and two-photon excited auto-fluorescence (TPEaF) imaging serve as prominent non-linear microscopy techniques with promising applications in cancer research, enabling label-free visualisation of non-centrosymmetric ordered structures (such as collagen) and fluorescence from autofluorophores (such as FAD/NADH) aiding investigations into the cell-matrix interface.

However, to date, the application of non-linear microscopy techniques for investigations into phenotypic collagen changes associated with OS is limited to few reports, particularly regarding clinical samples [9-11]. Thus, we have explored the potential of SHG microscopy to distinguish between normal human cortical bone and OS biopsies via label-free assessment of pathological alterations to the collagen matrix. We describe systematic optimisation of an imaging and analysis workflow to enable extraction of robust, fibre length-based parameters that have enabled accurate characterisation of healthy and pathological collagen signatures within clinical bone and OS biopsies across disease stage.

## Experimental Methods

### Samples

Decalcified human rib bone (n = 3) and human OS tissue (n = 3) biopsies were purchased from AMSBIO as paraffin-embedded arrays (each biopsy; 5 µm thick sections, 1-1.5 mm in diameter, product codes: BO244f, OS804D, OS208a). Tissue arrays were stored at 4°C and imaged at room temperature. Details on all biopsies used can be found in Supplementary T1.

### Second Harmonic Generation (SHG) / Two-Photon Excited Auto-Fluorescence (TPEaF) Imaging

SHG and TPEaF imaging was conducted using a custom-built, multi-photon imaging system (Supplementary Fig 1), as previously described [12] and MATLAB interface ScanImage 5.1 (Vidrio Technologies, USA). The system includes a 120-femtosecond pulsed laser (Mai Tai, Spectra-Physics) coupled through a galvanometric optical scanner, to an upright Leica DMRB microscope permitting simultaneous detection of SHG and TPEaF in separate channels. The laser was tuned at 800nm and passed through a quarter- (λ/4) and half- (λ/2) wave plate permitting control of laser polarisation. The beam is coupled through a galvanometric optical scanner and directed to the objective via a dichroic mirror (Semrock FF685-DI02-25×36). The emission signal is collected in the epi-detection and separated from the excitation beam by the same dichroic mirror and subsequently cleaned with a 694 nm cut-off short-pass filter. SHG was separated from other signals with a second dichroic mirror (Semrock, FF458-Di02) before being filtered (400 ± 20nm; Thorlabs, FB400-40) and focused onto a photomultiplier tube (PMT) (Hamamatsu, H10722-01). Similarly, TPEaF signal is filtered (520 ± 20 nm; Thorlabs, FBH520-40) and focused onto a separate PMT (Hamamatsu, H10722-20).

All SHG images were acquired using a 20x air objective with numerical aperture (NA) of 0.75 (Leica, Germany). The focal spot size (diffraction limited) of our microscope system is calculated as approximately 650 nm. A further 3x optical zoom (by limiting the scan angle) provided a FOV of ∼317 μm x 317 μm. Two image acquisitions were obtained over 8 ms per line at a resolution of 1024 × 1024 pixels. Laser power at the specimen was ∼30 mW. To overcome limited FOV, MATLAB was utilised for semi-automated acquisition of raster-scanned images in an 8 × 8 grid formation with 50% overlap between sequential images.

### Image Processing and Analysis

All image processing steps were completed using FIJI [13]. Intensity normalisation, noise removal and smoothing were conducted prior to semi-automated assembly of overlapping image tiles using the FIJI MIST plugin [14]. Extraction of collagen fibres from SHG images for quantification of fibre-length based parameters was completed using MATLAB-based CT-FIRE. CT-FIRE combines curvelet transform for image denoising and enhancement of fibre edges with the FIRE algorithm that enables extraction of individual fibres and descriptive metrics including fibre length [15]. A series of optimisation attempts was conducted to determine appropriate CT-FIRE input values for representative fibre extraction from SHG images, namely: a threshold value of 10, minimum fibre length of 50 and maximum fibre width of 15. Manipulation of these parameters can substantially influence the outcomes of fibre extraction, as demonstrated in Supplementary Fig 2.

## Statistical Analysis

Comparisons were undertaken using independent human biopsies per group, separated by biological sex (n = 3). Statistical analysis was performed on GraphPad Prism 9 (San Diego, CA, USA). A student’s t-test, one-way analysis of variance (ANOVA) or two-way ANOVA was conducted as appropriate, to compare the means of groups of data and the interaction of sex where appropriate. A Tukey’s post-hoc test was used for multiple comparisons. Data were presented as the mean ± standard deviation. Differences were considered to be statistically significant at P ≤ 0.05 (*P<0.05, **P<0.01, ***P<0.001).

The average top 50^th^ and lowest 50^th^ percentile values were calculated by averaging the longest 50% and shortest 50% of fibre lengths extracted from each individual biopsy respectively. The top and lowest 50^th^ percentiles were calculated to capture the specific effects of particularly long and particularly short collagen fibres, otherwise lost via overall averaging. Fibre length distribution graphs were created by classification of all fibre lengths (extracted by CT-FIRE) into 10 μm intervals between 10 - 400 μm and quantification of the percentage of total fibres within each length interval. Distributions are presented as the percentage of total fibres for normalisation across different FOV and biopsy sizes. Subsequent curve fitting analyses (GraphPad Prism 9) enabled quantification of several curve fit parameters describing the distribution profiles of fibre lengths. Specifically, a phase decay was modelled using the equation: Y = (Y0 – Plateau)*exp(-K*X) + Plateau. Here, Y0 is the Y value (percentage of total fibres) when X (fibre length) is zero. Plateau is the Y value at infinite X – thus reflecting the percentage of total fibres at the higher extremities of all fibre lengths exhibited. K is the rate constant, Tau is the time constant (reciprocal of K) and half-life is computed as In(“)/K, all indicators of curve steepness, and span is the difference between Y0 and plateau.

The number of fibres typically extracted from bone and OS biopsies using different FOV, from which collagen fibre length signatures were quantified, are detailed in Supplementary T2. All average collagen fibre parameters quantified herein are detailed in Supplementary T3 – 7.

## Results

Human bone and OS biopsies were imaged using SHG microscopy as sequential images with a 50% overlap and subsequently reconstructed in FIJI using the MIST plugin[14]. Final composite images up to 1400 µm x 1400 µm were analysed using MATLAB-based CT-FIRE for quantification of collagen fibre parameters enabling optimisation of the imaging methodology – image resolution and field of view (FOV) size – and comparison of human bone and OS collagen phenotypes. An overview of the experimental workflow utilised in these studies can be seen in Fig. 1.

**Fig 1.**
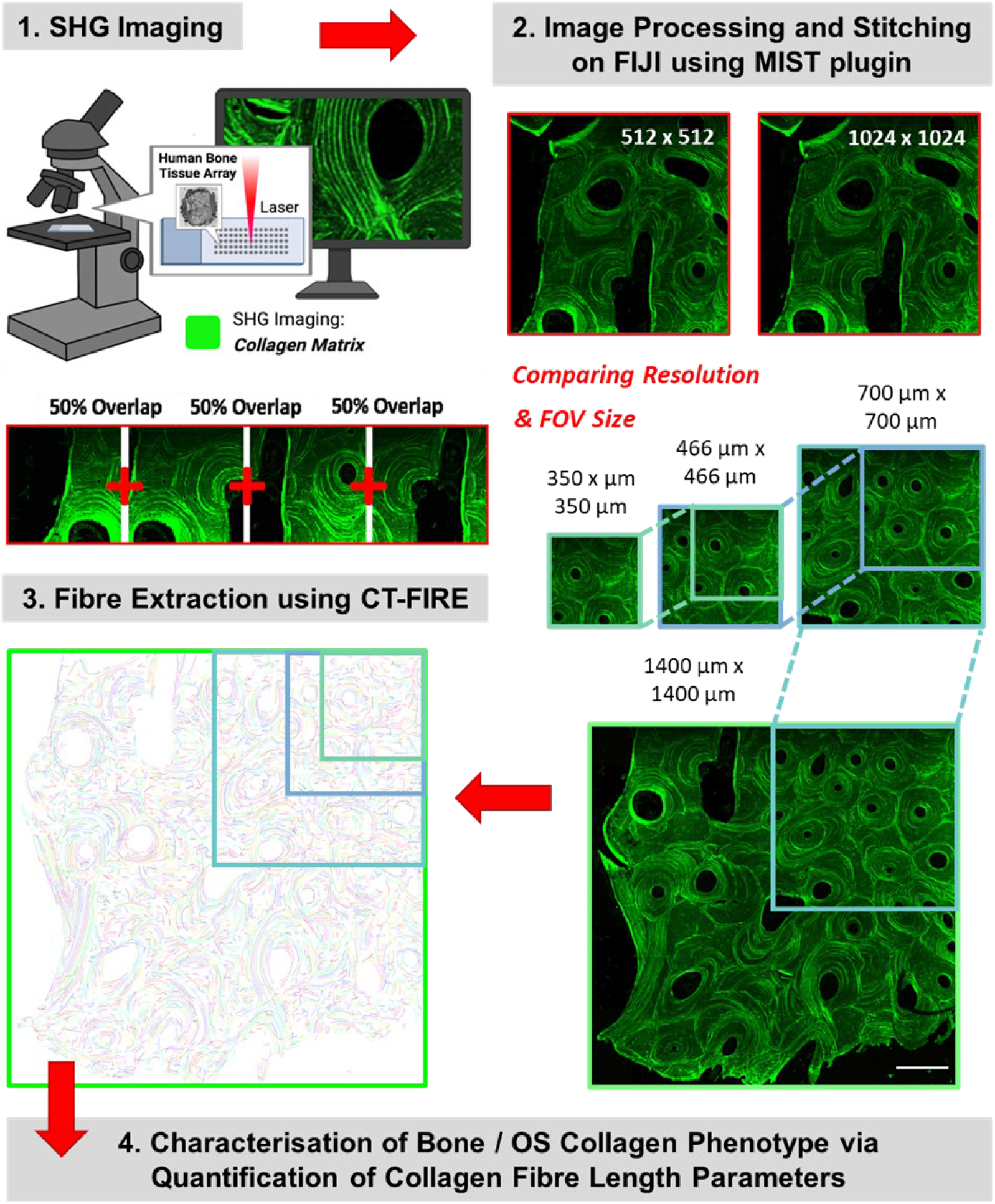
Experimental workflow for label-free characterisation of human (rib) bone and osteosarcoma biopsies. Human biopsies were imaged as 64 overlapping images with a 50% overlap (1) and subsequently reconstructed in FIJI using the MIST plugin (2). Final composite images (up to 1400 µm x 1400 µm) were analysed using MATLAB-based CT-FIRE (3) for quantification of collagen fibre parameters (4) enabling comparison of bone and osteosarcoma matrix properties.

### SHG image resolution significantly impacts collagen fibre analysis

Appropriate image resolution was determined by the acquisition of SHG/TPEaF images from three distinct human bone biopsies (n = 3), employing pixel dimensions of 512 × 512 and 1024 × 1024 (Fig. 2A, B). Subsequent comparison of collagen fibre parameters obtained by CT-FIRE demonstrated the significant impact of image resolution on collagen fibre analysis. All average values quantified are detailed in Supplementary T3. Average fibre density significantly decreased (p < 0.001) in 512 × 512 pixel images compared to 1024 × 1024 pixel images (Fig. 2C). Average fibre straightness also decreased (p < 0.05) in 512 × 512 pixel images compared to 1024 × 1024 pixel images (Fig. 2D). Conversely, average fibre width significantly increased (p < 0.001) in 512 × 512 pixel images compared to 1024 × 1024 pixel images (Fig. 2E). Average fibre length also significantly increased (p < 0.01) in 512 × 512 pixel compared to 1024 × 1024 pixel images (Fig. 2F). This divergence was observed for shorter fibres (lowest 50^th^ percentile of fibre lengths) which significantly increased (p < 0.01) in 512 × 512 pixel images compared to 1024 × 1024 pixel images (Fig. 2G), as well as for longer fibres (top 50^th^ percentile of fibre lengths) which also demonstrated a significant increase (p = 0.01) in 512 × 512 pixel images compared with 1024 × 1024 pixel images (Fig. 2H).

**Fig 2.**
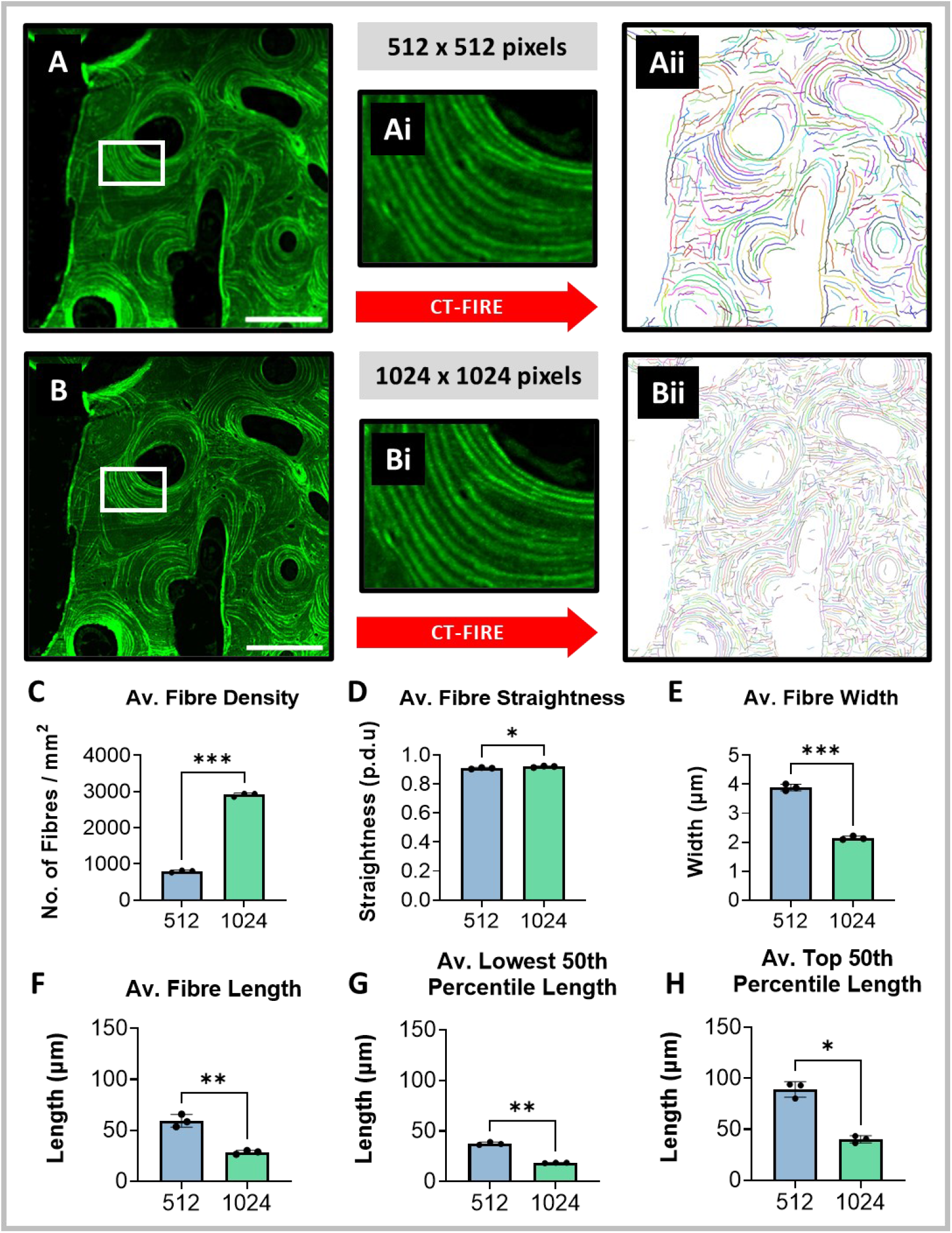
Significant differences in collagen fibre parameters acquired from 512 × 512 pixel and 1024 × 1024 SHG images of human bone tissue. 512 × 512 pixel (Ai, Aii) and 1024 × 1024 pixel (Bi, Bii) SHG images were acquired of human bone for comparison of collagen fibre using CT-FIRE (Aiii, Biii). Collagen parameters quantified included average: fibre density (C), fibre straightness (D), fibre length (E), fibre width (F), top 50^th^ percentile length (G) and lowest 50^th^ percentile length (H). Scale bars correspond to 200 µm. Data represents the mean of three independent samples (n = 3) with error bars denoting ± SD (*P<0.05, **P<0.01, ***P<0.001).

Notably, despite clear visibility of the collagen matrix in both 512 × 512 and 1024 × 1024 pixel images, every collagen parameter quantified exhibited significant differences between the two image resolutions. These findings demonstrate the significant information loss that may accompany diminished pixel resolution, and the importance of microscopic optimisation. This finding aligns with the Nyquist theorem, which states that the sampling rate must equal at least twice the highest frequency present in the signal, that is, the pixel size must be at least 2-3 times smaller than the feature that is being resolved. Our resolving power is approximately 650 nm while collagen fibril width has been reported to extend up to 500 nm, demonstrated by both electron microscopy and SHG imaging [12, 16, 17]. 1024 × 1024 pixel images result in a pixel size of 310 nm, whilst 512 × 512 pixel images yield a less favourable pixel size of 619 nm. Given our resolving power, a pixel size of 310 nm is considerably better suited to image and discern smaller collagen fibres. Consequently, all subsequent imaging was carried out at a resolution of 1024 × 1024 pixels, despite the notable 4x increase in time requisite for image acquisition.

### Determining optimal field of view size on collagen fibre length distributions from normal bone and OS biopsies

To ensure accurate characterisation of normal and pathological bone collagen phenotypes, the influence of image FOV size on quantification of collagen parameters was assessed. SHG images were acquired of bone and stage IIB OS biopsies (n = 3) and processed to provide composite images of four different FOV sizes: 350 µm x 350 µm, 466 µm x 466 µm, 700 µm x 700 µm, 1400 µm x 1400 µm (Fig. 3A, Fig. 4A). CT-FIRE enabled fibre extraction (Fig. 3B, Fig. 4B) and quantification of fibre lengths – detailed in supplementary table 4. In normal bone biopsies, no significant difference was observed in average fibre length values extracted across all FOV sizes examined (Fig. 3Ci). Significant differences were present, however, between the longest fibre lengths extracted from different FOV sized SHG images of human bone. The top 50^th^ percentile of fibres significantly increased in length with increasing FOV size until 700 µm x 700 µm (Fig. 3Cii). Plotting the distribution of all collagen fibre lengths extracted from a human bone biopsy per FOV size further highlighted the significant impact of FOV size on accurately capturing the longest collagen fibres present within human bone biopsies (Fig. 3D).

**Fig 3.**
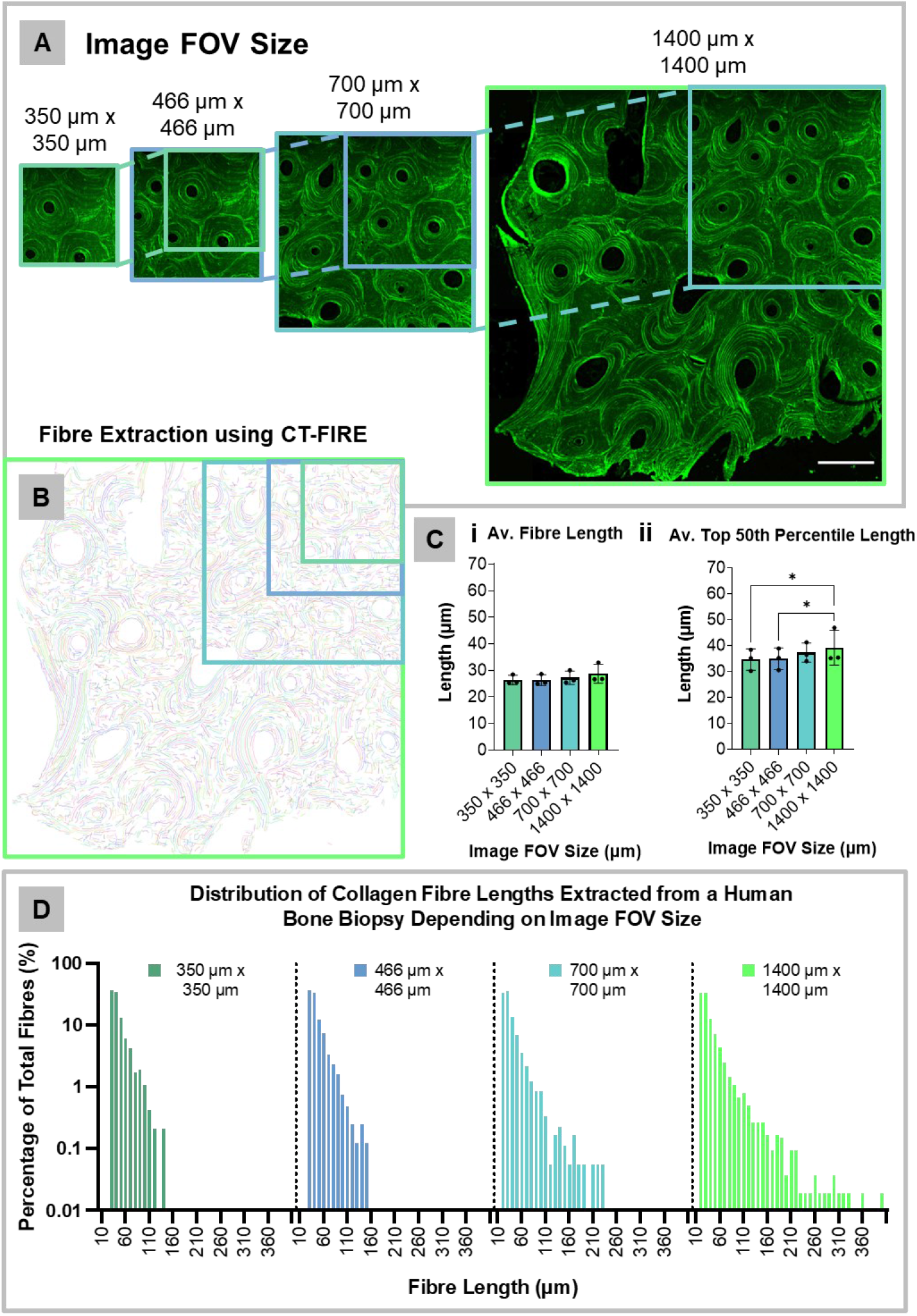
Larger field of view (FOV) size improves extraction of collagen fibre lengths from normal human bone biopsies. A) SHG images of different FOV sizes as indicated. B) Collagen fibre extraction by CT-FIRE. C) Quantification of average fibre length (µm; Ci) and average top 50^th^ percentile length (µm: Cii) for each FOV size. Data represents the mean of three independent biopsies (n = 3) ± SD. Significance was assessed using one-way ANOVA (P<0.05 *). D) Distribution of all collagen fibre lengths presented as a percentage of total fibres extracted from a single human bone biopsy across each FOV size.

**Fig 4.**
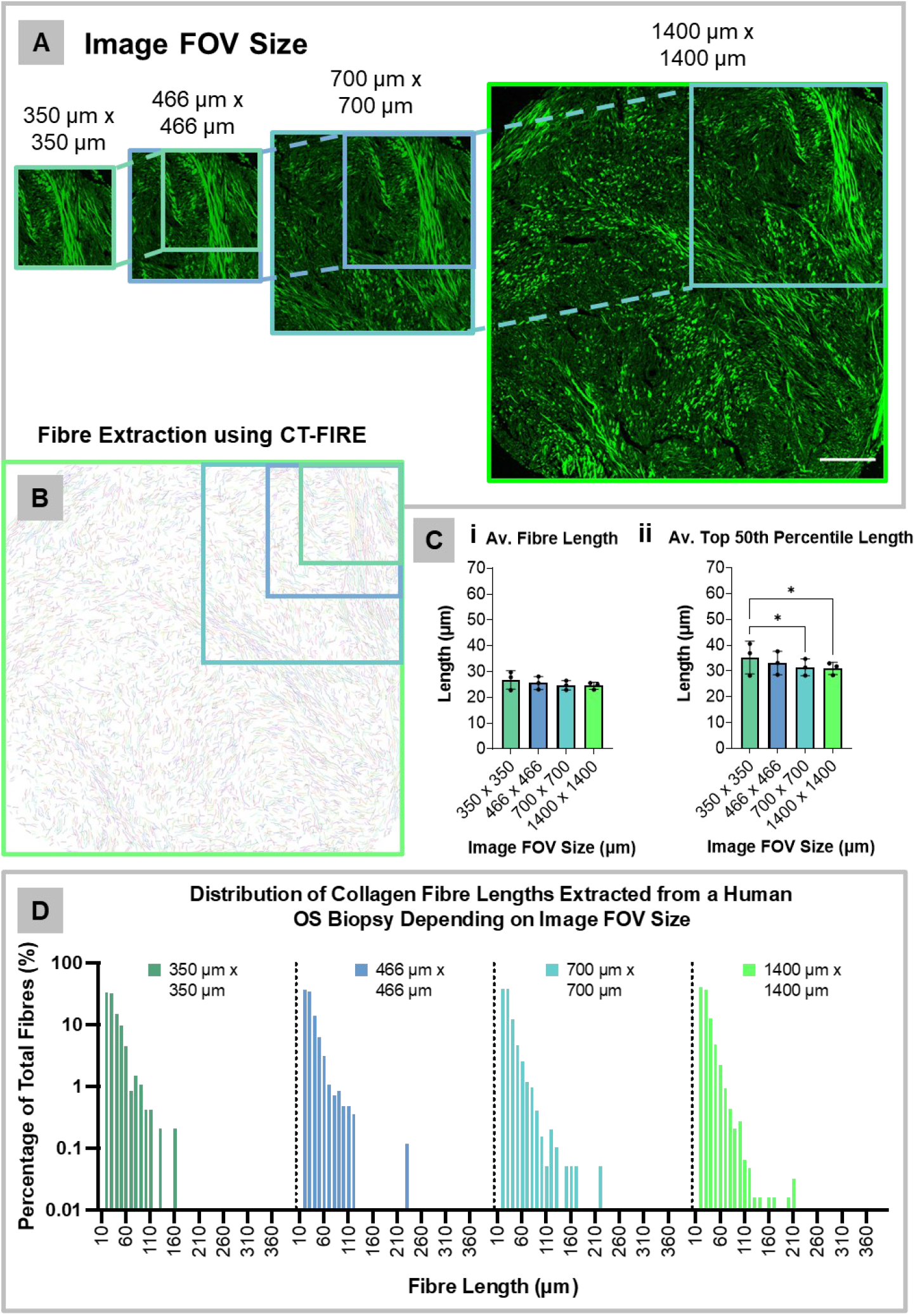
Larger field of view (FOV) improves extraction of collagen fibre lengths from osteosarcoma (stage IIB) biopsies. A) SHG images of different FOV sizes as indicated. B) Collagen fibre extraction by CT-FIRE. C) Quantification of average fibre length (µm; Ci) and average top 50^th^ percentile length (µm: Cii) for each FOV size. Data represents the mean of three independent biopsies (n = 3) ± SD. Significance was assessed using one-way ANOVA (P<0.05*). D) Distribution of all collagen fibre lengths presented as a percentage of total fibres extracted from a single human OS biopsy across each FOV size.

In line with this, no significant differences were observed in the average fibre lengths extracted from different FOV sized images of OS (stage IIB) biopsies (Fig. 4Ci). Again, however, significant differences were detected in the top 50^th^ percentile of fibre lengths extracted from the different FOV sized images. In contrast to normal bone, the top 50^th^ percentile of fibres in OS biopsies significantly decreased in length with increasing FOV size (Fig. 4Cii). The total distribution of collagen fibre lengths extracted from a human OS biopsy at each FOV size also illustrated the impact of image FOV size on collagen fibre analysis (Fig. 4D). Such findings potentially reflect the high heterogeneity of OS biopsy collagen, which is more accurately represented with a larger FOV size.

### Comparative SHG / TPEaF imaging of human bone and osteosarcoma

Label-free images of human bone (Fig. 5A female, 5B male) and stage IIB OS (Fig. 5C female, 5D male) biopsies were acquired by stitching 64 overlapping images (317 µm x 317 µm) representing a total area of 1400 µm x 1400 µm. Collagen fibres visualised by SHG imaging are represented in green, with TPEaF represented in blue and images overlaid. In normal bone, collagen fibres were clearly identified forming organised circular layers around the Haversian canals containing blood vessels (blue; Fig 5Ai/Bi). SHG imaging highlighted distinct collagen patterning in OS soft tissue biopsies, with the collagen matrix displaying a highly disorganised and poorly structured composition with large amounts of collagen encompassing cell nuclei (Fig 5Ci/Di).

**Fig 5.**
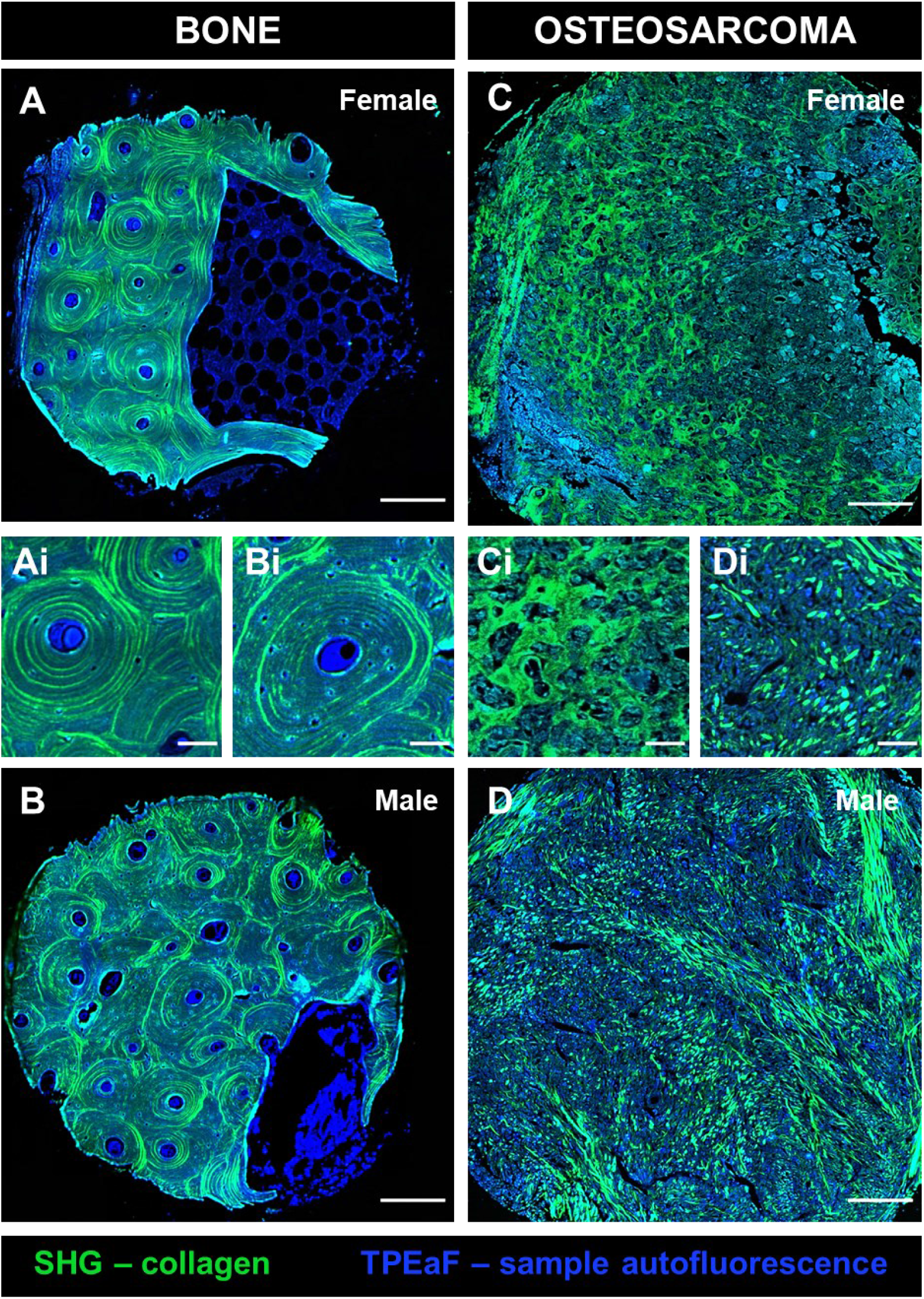
Type 1 collagen distribution in normal bone and osteosarcoma biopsies acquired by SHG imaging. SHG (green) and TPEaF (blue) images were acquired of human bone and OS (stage IIB) tissue and overlaid for visualisation of collagen matrix interface. Tiling of 64 overlapping images enabled generation of the composite images visualised above. Scale bars; 200µm (A-D), 50µm (Ai-Di).

### Collagen fibre lengths are reduced in osteosarcoma

To assess whether image FOV size impacts comparisons between human bone and OS collagen, analyses were undertaken on SHG images across the four FOV sizes generated (n = 3). The average length and top 50^th^ percentile length of collagen fibres exhibited in normal bone and OS were compared per FOV size, with biopsies separated by sex (Fig. 6). Significant differences between normal bone and OS (stage IIB) collagen were identified across both female and male comparisons, with OS biopsies exhibiting reduced average and top 50^th^ percentile fibre lengths compared to normal bone. Notably, however, FOV size had a significant influence on this ability to identify distinctions between normal bone and OS collagen. Significant differences were only identified in male comparisons when analysing SHG images with the larger FOV sizes of 700 µm x 700 µm and 1400 µm x 1400 µm. OS-associated reductions in collagen fibre lengths were concealed when using the smaller FOV sizes of 350 µm x 350 µm and 466 µm x 466 µm, highlighting the importance of FOV size optimisation for accurately phenotyping clinical bone collagen. Though distinctions between female bone and OS collagen were permissible using the smaller FOV sizes, differences were augmented when using both the larger FOV sizes of 700 µm x 700 µm and 1400 µm x 1400 µm. All average and top 50^th^ percentile fibre lengths quantified are detailed in Supplementary T5.

**Fig 6.**
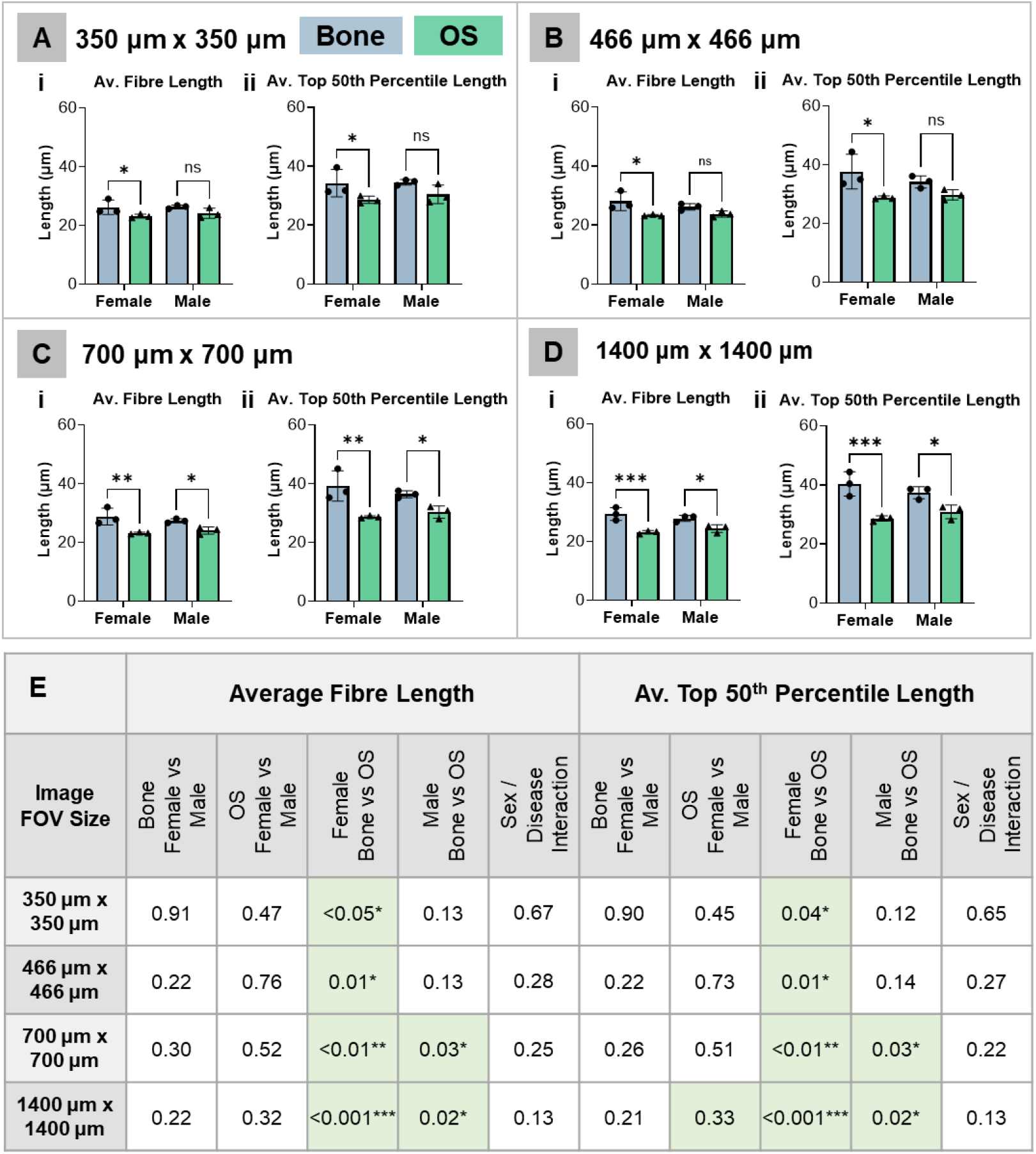
Reduced collagen fibre lengths in osteosarcoma evident only across larger FOV. Average fibre length (i) and average top 50^th^ percentile length (ii) extracted from SHG images of sex-matched normal human bone and OS biopsies. Fibre analysis was undertaken on images with a FOV size of 350 µm x 350 µm, (A) 466 µm x 466 µm (B) 700 µm x 700 µm (C) and 1400 µm x 1400 µm (D). Table summarising probability values for comparisons drawn and sex/disease interaction (E). Data represents the mean of three independent samples (n = 3) with error bars denoting ± SD. Significance was assessed using two-way ANOVA (*P<0.05, **P<0.01, ***P<0.001).

### Mapping collagen fibre length distributions in across human bone and osteosarcoma biopsies

To further explore novel metrics for OS tissue identification based on collagen fibre lengths, curve fitting analysis was applied to total fibre length distributions of individual bone and OS (stage IIB) biopsies (n = 3), using the maximum FOV of 1400 µm x 1400 µm (Fig. 7). Curve fitting analyses enabled quantification of several parameters describing the overall distribution pattern of collagen fibre lengths exhibited within individual biopsies: Y0, Plateau, K, Half-Life, Tau and Span (Fig. 7A). These parameters provide a more holistic overview of collagen fibre length phenotypes and permitted distinction between normal bone and OS biopsies (Fig. 7B, C). Further detail on this analysis and the parameters acquired can be found in the methodology section.

**Fig 7:**
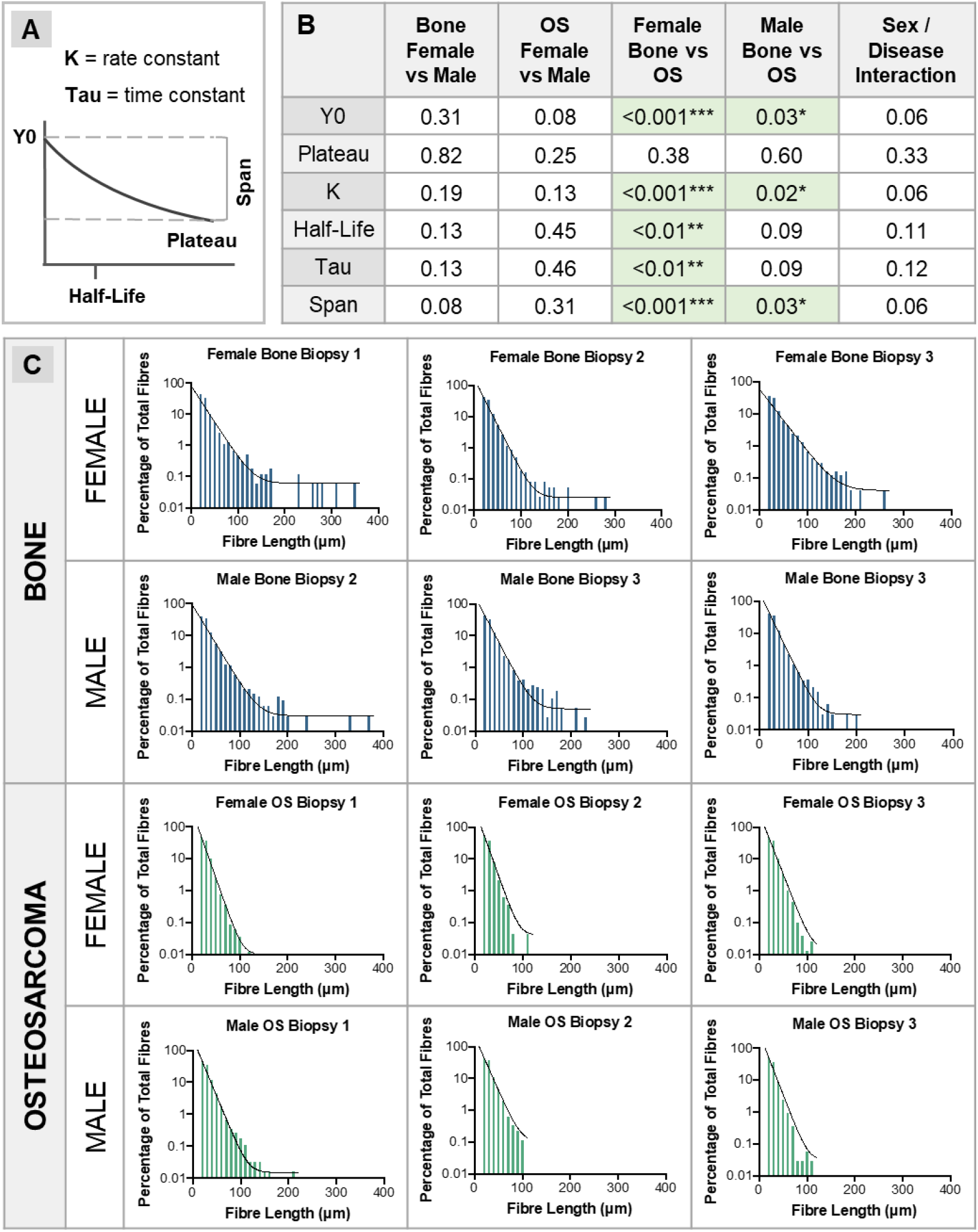
Distribution of collagen fibre lengths across normal bone and osteosarcoma biopsies. Curve fitting analysis on fibre length distributions enabled quantification of a range of curve fit parameters (A). Probability values for comparisons drawn and interaction between sex and disease (B). Distributions of fibre lengths extracted from individual bone and stage IIB OS biopsies from both sexes (C). Data represents the mean of three independent samples (n = 3). Significance was assessed using two-way ANOVA (*p<0.05, <0.01**, p<0.001***).

The curve fit parameters Y0, K (rate constant) and Span all demonstrated significant differences between bone and OS in both female and male comparisons. Half-Life and Tau both demonstrated a significant difference between bone and OS in female comparisons but not male. Of the various curve fit parameters, plateau was the only parameter with no ability to differentiate between normal bone and OS. The significant differences exhibited in the fibre length distribution patterns between normal bone and OS validated the diagnostic potential of these novel curve fitting metrics. In contrast, no significant interaction between sex and disease was reported for any of the curve fitting parameters explored following two-way ANOVA.

### Characterisation of collagen fibre signatures with OS disease stage

Following initial exploration of our optimised SHG methodology in revealing cancer-associated changes to bone collagen signatures in stage IIB OS, we further investigated the potential of this method in examining OS progression – from stage IB to IVB tumours. Label-free SHG / TPEaF images were acquired of male: rib bone (Fig. 5B), stage IB OS (Fig. 8A), stage IIB OS (Fig. 8B), stage III OS (Fig. 8C) and stage IVB OS (Fig. 8D), biopsies and overlaid for visualisation of the matrix interface. A total of 30 biopsies were examined for this investigation – six biopsies per group – with repeats from two (stage III and IVB OS) or three (bone, stage IB and IIB OS) independent patients depending on tissue availability. Visually, OS tissue exhibits a vastly dissimilar ECM phenotype compared to normal bone.

**Fig 8.**
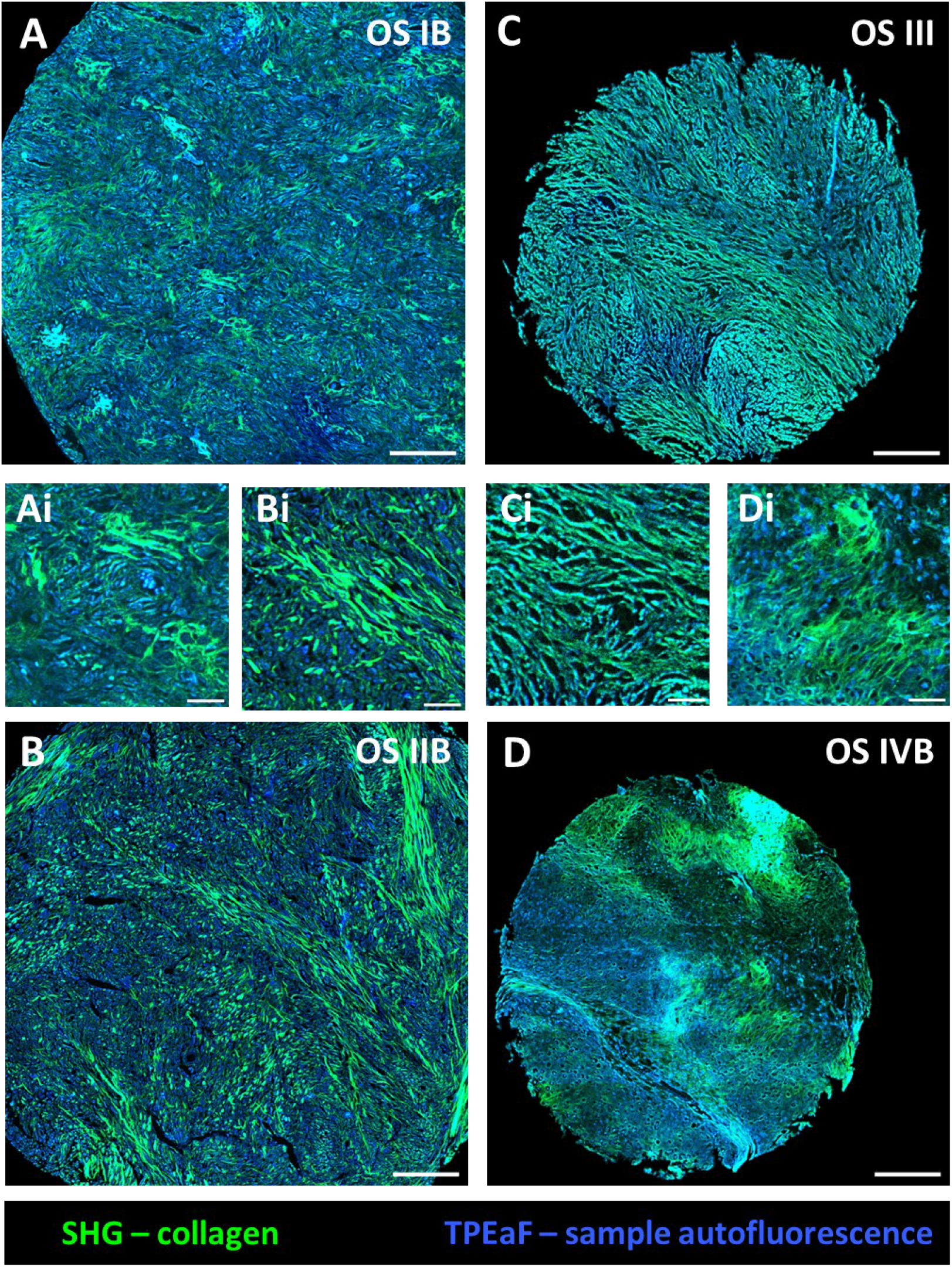
The collagen matrix of osteosarcoma biopsies with disease stage visualised SHG/TPEaF imaging. SHG (green) and TPEaF (blue) images were acquired of male: stage IB OS (A), stage IIB OS(B), stage III OS (C) and stage IVB OS (D) biopsies and overlaid for visualisation of the collagen matrix interface. Entire biopsies were visualised by stitching 64 overlapping images. Scale bars correspond to 200 µm (A-D), 40 µm (Ai-Di).

Quantification of collagen fibre lengths highlighted a reduction in collagen fibre lengths with cancer progression. When investigating across progressive OS stages, normal bone biopsies could be differentiated from advanced OS tumours (stage III, IVB), but not from early-stage OS tumours (stage IB, IIB). Normal bone exhibited a significantly increased average fibre length compared to stage III OS (p = 0.04) and stage IVB OS (p < 0.05) (Fig. 9I). This suggests OS-associated reductions in collagen fibre lengths are exacerbated in advanced stages of the disease, coinciding with the presence of metastasis[18]. Earlier-stage OS biopsies also demonstrated reductions in average fibre length but did not reach significance. We hypothesise that significance would be met with a larger sample size – as evidenced by our bone-stage IIB OS only comparisons. Analogously, the top 50^th^ percentile of fibres also demonstrated a significant length reduction in stage III (p = 0.04) and stage IVB (p = 0.04) biopsies compared to normal bone (Fig. 9K). Conversely, the lowest 50^th^ percentile did not show any significant difference between normal bone and any of the OS biopsies investigate (Fig. 9J).

**Fig 9.**
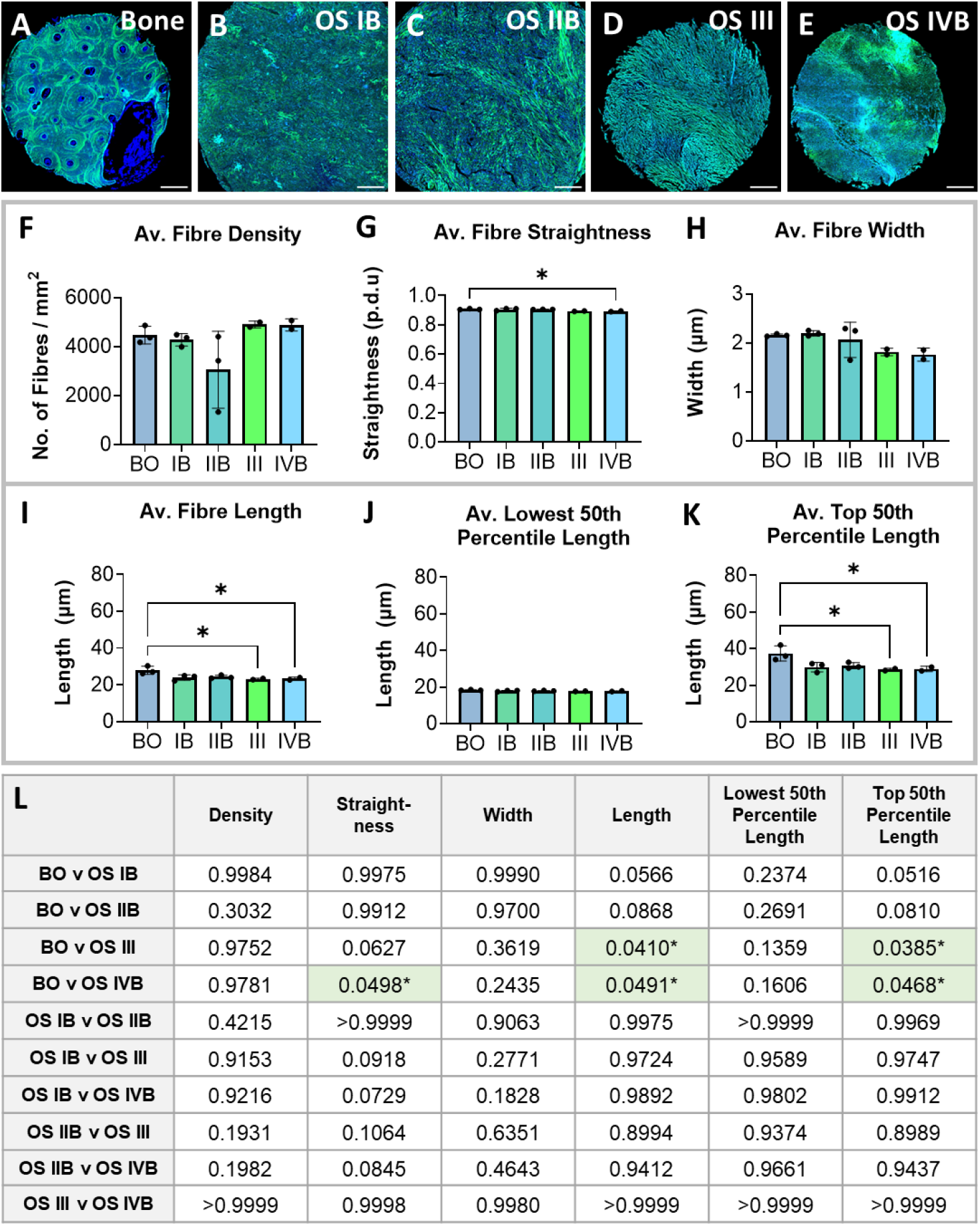
Collagen fibre parameters acquired from SHG images of bone and osteosarcoma biopsies with increasing cancer stage. SHG/TPEaF images were acquired of male: bone (A), stage IB (B), stage IIB (C), stage III (D) and stage IVB (E) OS biopsies. CT-FIRE analysis of SHG images enabled comparison of collagen fibre metrics. Entire biopsies were visualised by stitching 64 overlapping images. Average fibre density (F), fibre straightness (G), fibre width (H), fibre length (I), length of shortest fibres (J) and length of longest fibres (K) were quantified. Data represents the mean of six biopsies in total, either from three (bone, IB and IIB OS, n = 3) or two (III, IVB OS, n = 3) independent patients dependent on tissue availability. Error bars denote ± SD. Significance was assessed using one-way ANOVA (L) (P<0.05*). Scale bars correspond to 200 µm.

Other non-length-based collagen fibre parameters were also quantified in the bid to determine phenotypic differences between normal bone and OS collagen. Average fibre density (Fig. 9F) and average fibre width (Fig. 9H) demonstrated no significant difference between normal bone and any of the OS biopsies examined. However, average fibre width demonstrated an obvious – albeit insignificant – trend of decline with increased malignancy stage. Thus, with a larger sample size it is hypothesised that collagen fibre width may be another quantitative parameter with diagnostic and prognostic promise. Lastly, average fibre straightness, also exhibited a consistent decrease with advancement of OS (Fig. 9G). However, a statistically significant reduction was only observed in bone – stage IVB OS biopsy comparisons with this sample size (p = 0.05). Again, these results suggest that pathological collagen phenotypes accommodating OS may intensify with disease progression and demonstrate the potential of SHG imaging in monitoring such manifestations. All collagen fibre parameters quantified are detailed in Supplementary T7.

## Discussion

The current studies provide evidence for label-free detection of OS phenotypes from clinical biopsies through systematic characterisation of aberrant collagen fibres visualised by SHG imaging. The alteration of OS-collagen fibres observed herein reflects findings from previous studies that have examined OS using non-linear microscopy [9-11]. Our studies have shown that appropriate image resolution and field of view size are critical for accurately assessing collagen fibre length as a diagnostic biomarker of human bone pathology. Specifically, a FOV size of 700 µm × 700 µm or greater was required to accurately phenotype human bone and OS biopsies from SHG images. Smaller FOVs resulted in a loss of information, particularly regarding the longest fibres present, compromising phenotype and diagnostic analyses. In OS biopsies, fibre length decreased with increasing FOV, likely reflecting the high heterogeneity of the tumour matrix requiring a large FOV for accurate characterisation. Fibre length distribution assessments, achieved via curve fitting analyses, revealed novel length-based metrics that significantly differentiated normal bone from OS biopsies. Furthermore, SHG imaging of OS biopsies across clinical stages (IB–IVB) demonstrated that collagen abnormalities become more pronounced with disease progression, highlighting the diagnostic value of this workflow.

Utilising SHG imaging to determine collagen fibre-length based metrics remains relatively underexplored as a phenotypic and potential diagnostic parameter in marked contrast to metrics such as collagen density and fibre organisation, which include collagen orientation and alignment [9, 19-23]. From the studies that have considered collagen fibre length, fibre lengths are typically averaged across the sample on analysis [24, 25]. This method may lack the sensitivity required to capture subtle disparities between tissue types or may introduce disparities based on FOV size. Historically, there is a large discrepancy in the range of FOV sizes used for SHG examination of human tissue collagen, ranging from <200 µm x 200 µm to >1000 µm x 1000 µm [19, 21, 23, 25-27]. Though stitching of SHG images enabling visualisation of whole biopsy cores has been reported, collagen fibre analysis and quantification is typically performed on individual SHG images acquired or smaller ROIs [22, 27]. Moreover, despite the significant influence of microscopic parameters such as FOV size on research findings, optimisation of such variables is typically not reported.

It is important to note that no significant differences were identified in female-male comparisons of bone biopsies or OS biopsies, across the collagen fibre parameters investigated. However, the magnitude of divergence between normal and pathological collagen varied between female versus male comparisons – suggesting OS collagen and ECM heterogeneity may exhibit sexual dimorphism. Notably, men demonstrate higher OS incidence [28] and notably poorer survival of OS than women [29], which could link to complexities underlying sexually dimorphic manifestations of the disease. Further investigations into the mechanisms driving OS progression via aberrant ECM alterations, and the influence of sex, could inform personalised therapies and improve patient outcomes.

The non-invasive nature of label-free modalities such as SHG microscopy present valuable, non-destructive alternatives to study, examine and potentially diagnose disease pathology. The practicality of employing such techniques in clinical settings has been validated by near real-time, intraoperative diagnosis of brain tumours [30].

Label-free, non-linear imaging combined with deep learning enabled precise delineation of tumour boundaries and automated diagnosis in the operating room. Diagnoses were achieved at a comparable accuracy to conventional pathologist-based interpretation of histological images but considerably faster at <150s versus >20 min [30]. Such results demonstrate how cancer diagnosis may be streamlined and completed intraoperatively, avoiding the need for separate pathological analysis laboratories [30]. In addition to improving current diagnosis pathways, the ability to delineate tumour boundaries more precisely will enhance the accuracy of tumour resection and thus reduce the risk of disease recurrence [30, 31]. Such improvements hold particular significance for aggressive cancers, such as high-grade OS, where prognosis is bleak, and the threat of disease relapse persists. Thus, continuous improvements to analytical methods enabling rapid, automated analysis are key to the prospects of label-free microscopic diagnosis in clinical settings.

Our findings highlight the potential for robust and systematic quantitative assessments of human bone collagen, with optimisation of imaging parameters and analysis methods critical for accurate interpretation and potential clinical diagnosis. Collagen fibre length-based metrics particularly, show promise as novel diagnostic signatures of OS and suggest therapeutic relevance. Further standardisation of collagen quantification across tissues and disease states could enable broader application of SHG imaging in cancer diagnostics clinically.

## Acknowledgements

This studentship was funded by Hannah’s Willberry Wonder Pony, UK (registered charity number: 1166416) and the University of Southampton, UK. Contribution from Engineering and Physical Sciences Research Council (EPSRC) grant (EP/T020997/1) is also acknowledged for supporting equipment and microscopy related aspects of this work.

