## Supplementary for "Characterising Type I Collagen Architecture in Human Bone and Osteosarcoma with Second Harmonic Generation Microscopy"

**Supplementary Table 1.** Details of human bone and osteosarcoma biopsies used in these studies. Colour-coding details biopsies used for more than one investigation.

| Purpose | Tissue | Sex | Age | Site | No. of Biopsies |
|---|---|---|---|---|---|
| Image Optimisation | Bone | Male | 56 | Rib | 1 |
| | | | 57 | Rib | 1 |
| | | | 58 | Rib | 1 |
| Validation of Optimised Protocol | Bone | Female | 62 | Rib | 1 |
| | | | 66 | Rib | 1 |
| | | | 66 | Rib | 1 |
| | | Male | 62 | Rib | 1 |
| | | | 66 | Rib | 1 |
| | | | 67 | Rib | 1 |
| | OS (IIB) | Female | 12 | Femur | 1 |
| | | | 14 | Femur | 1 |
| | | | 16 | Femur | 1 |
| | | Male | 12 | Femur | 1 |
| | | | 14 | Femur | 1 |
| | | | 16 | Femur | 1 |
| Distinction of Normal Bone from Different Stages of OS | Bone | Male | 62 | Rib | 2 |
| | | | 66 | Rib | 2 |
| | | | 67 | Rib | 2 |
| | OS (IB) | Male | 18 | Femur | 2 |
| | | | 18 | Humerus | 2 |
| | | | 21 | Femur | 2 |
| | OS (IIB) | Male | 12 | Femur | 2 |
| | | | 14 | Femur | 2 |
| | | | 16 | Femur | 2 |
| | OS (III) | Male | 15 | Humerus | 3 |
| | | | 46 | Humerus | 3 |
| | OS (IVB) | Male | 17 | Humerus | 3 |
| | | | 28 | Leg | 3 |

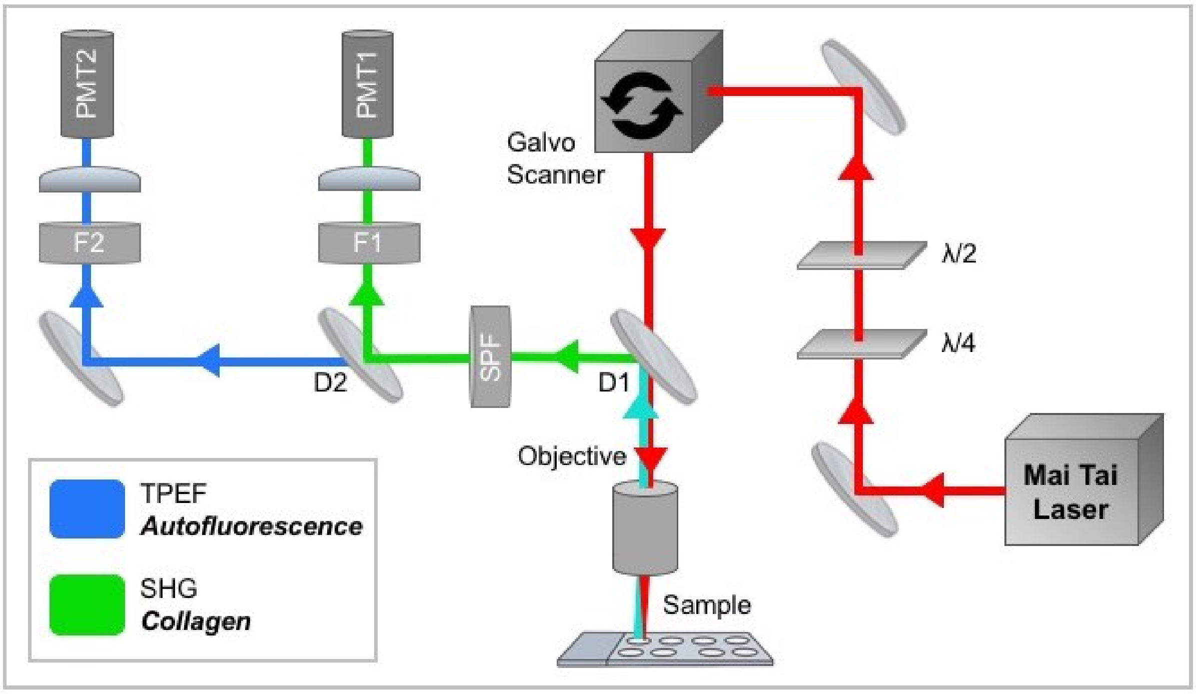


**Supplementary Fig 1. Multi-photon imaging system used to acquire label-free SHG images.** The femtosecond pulsed Titanium (TI): Sapphire laser was tuned at 800 nm (red) and passed through a quarter- (λ/4) and half- (λ/2) wave plate permitting control of laser polarisation. The beam is coupled through a galvanometric optical scanner and dichroic mirror (D1) enabling separation of excitation and emission. Signal is cleaned with a short-pass filter (SPF) and SHG (green) is separated with a second dichroic mirror (D2) before being filtered (F1) and focused onto a photomultiplier tube (PMT1) detector. TPEaF (blue) is similarly filtered (F2) and focused onto PMT2.

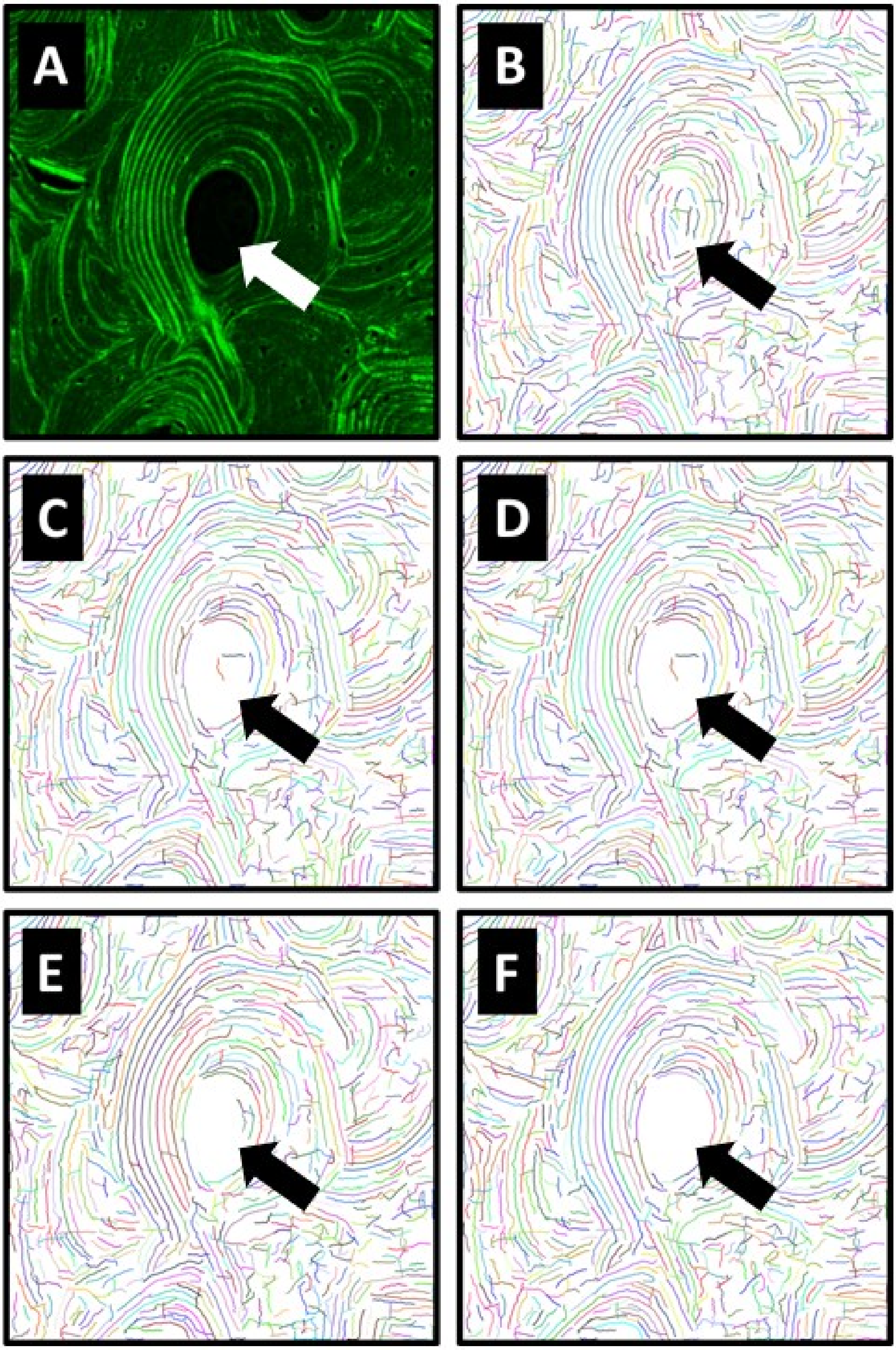


**Supplementary Fig 2. Example differences in fibre extraction from SHG images depending on input settings of CT-FIRE.** For CT-FIRE analysis of SHG images, a threshold value, minimum fibre length and maximum fibre width must be manually selected. The difference in fibre extraction from SHG images of human bone (A), using different threshold values of 2 (B), 4 (C), 6 (D), 8 (E), 10 (F) is shown.

**Supplementary Table 2.** Details on number of fibres extracted from human bone and osteosarcoma biopsies during methodology optimisation. Data represents the mean of three independent samples with error bars denoting ± standard deviation.

| Image FOV Size | Bone | | OS | |
|---|---|---|---|---|
| | No. of fibres extracted | No. of fibres per mm$^2$ of tissue | No. of fibres extracted | No. of fibres per mm$^2$ of tissue |
| **350 µm x 350 µm** | 427 ± 96.72 | 3733 ± 610.30 | 321 ± 210.60 | 2623 ± 1719.00 |
| **466 µm x 466 µm** | 770 ± 86.17 | 3711 ± 299.70 | 608 ± 315.70 | 2798 ± 1454.00 |
| **700 µm x 700 µm** | 1715 ± 131.60 | 3597 ± 261.00 | 1394 ± 663.70 | 2845 ± 1355.00 |
| **1400 µm x 1400 µm** | 4938 ± 799.20 | 3405 ± 257.30 | 3822 ± 2262 | 2011 ± 1166.00 |

**Supplementary Table 3:** Average collagen fibre parameters extracted from SHG images of human bone using two image resolutions (512 x 512 pixels versus 1024 x 1024 pixels) during methodology optimisation. Data represents the mean of three independent samples ± standard deviation.

| Collagen Fibre Parameter | SHG Image Pixel Resolution | |
|---|---|---|
| | 512 x 512 | 1024 x 1024 |
| Density (fibres / mm²) | 797 ± 31.05 | 2911 ± 51.10 |
| Straightness (p.d.u) | 0.91 ± < 0.01 | 0.92 ± < 0.01 |
| Width (µm) | 3.89 ± 0.12 | 2.15 ± 0.06 |
| Length (µm) | 59.31 ± 6.19 | 28.53 ± 1.82 |
| Lowest 50th Percentile Length (µm) | 37.55 ± 1.58 | 18.57 ± 0.26 |
| Top 50$^{th}$ Percentile Length (µm) | 89.03 ± 7.71 | 40.16 ± 2.03 |

**Supplementary Table 4:** Average collagen fibre length and top 50th percentile length extracted from SHG images of human bone and osteosarcoma biopsies per FOV size during methodology optimisation. Data represents the mean of three independent samples ± standard deviation.

| Tissue Type | Collagen Fibre Parameter | FOV Size | | | |
|---|---|---|---|---|---|
| | | 350 µm x 350 µm | 466 µm x 466 µm | 700 µm x 700 µm | 1400 µm x 1400 µm |
| Bone | Average Length | 26.44 ± 1.75 µm | 26.34 ± 2.01 µm | 27.25 ± 2.47 µm | 28.78 ± 3.58 µm |
| | Av. Top 50th Percentile Length | 34.63 ± 4.13 µm | 34.92 ± 2.42 µm | 37.30 ± 3.72 µm | 39.18 ± 6.72 µm |
| OS (IIB) | Average Length | 26.77 ± 3.57 µm | 25.56 ± 2.50 µm | 24.67 ± 1.79 µm | 24.51 ± 1.35 µm |
| | Av. Top 50th Percentile Length | 35.20 ± 6.34 µm | 33.09 ± 4.53 µm | 31.41 ± 3.22 µm | 31.03 ± 2.38 µm |

**Supplementary Table 5:** Average collagen fibre length and top 50th percentile length extracted from SHG images of human bone and osteosarcoma biopsies per FOV size, separated by sex. For determination of the influence of image FOV size on the ability to discriminate OS from normal bone using collagen fibre analysis. Data represents the mean of three independent samples ± standard deviation.

| Sex | FOV Size | Average Fibre Length (µm) | | Average Top 50th Percentile Length (µm) | |
|---|---|---|---|---|---|
| | | Bone | OS (IIB) | Bone | OS (IIB) |
| Female | 350 µm x 350 µm | 26.21 ± 2.48 | 23.22 ± 0.65 | 34.27 ± 4.64 | 28.60 ± 1.22 |
| | 466 µm x 466 µm | 28.11 ± 3.15 | 23.35 ± 0.31 | 37.69 ± 5.89 | 28.83 ± 0.58 |
| | 700 µm x 700 µm | 28.84 ± 2.85 | 23.24 ± 0.29 | 39.24 ± 5.18 | 28.73 ± 0.39 |
| | 1400 µm x 1400 µm | 29.39 ± 2.17 | 23.18 ± 0.56 | 40.29 ± 4.12 | 28.62 ± 0.92 |
| Male | 350 µm x 350 µm | 26.37 ± 0.59 | 24.19 ± 1.74 | 34.58 ± 0.92 | 30.49 ± 3.17 |
| | 466 µm x 466 µm | 26.21 ± 1.13 | 23.80 ± 1.02 | 34.18 ± 2.06 | 29.77 ± 1.74 |
| | 700 µm x 700 µm | 27.41 ± 0.70 | 24.10 ± 1.20 | 36.42 ± 1.21 | 30.37 ± 2.09 |
| | 1400 µm x 1400 µm | 27.87 ± 1.05 | 24.39 ± 1.32 | 37.35 ± 2.04 | 30.85 ± 2.32 |

**Supplementary Table 6:** Average curve fit parameters quantified following curve fitting analysis on the total distribution of fibre lengths extracted from bone and osteosarcoma biopsies, separated by sex. Data represents the mean of three independent samples ± standard deviation.

| Curve Fit Parameter | Female | | Male | |
|---|---|---|---|---|
| | Bone | OS (IIB) | Bone | OS (IIB) |
| **Y0** | 101.40 ± 55.59 | 332.23 ± 45.16 | 144.05 ± 41.55 | 251.13 ± 51.15 |
| **Plateau** | 0.04 ± 0.02 | 0.02 ± 0.02 | 0.04 ± 0.01 | 0.05 ± 0.05 |
| **K** | 0.06 ± 0.01 | 0.10 ± < 0.01 | 0.07 ± < 0.01 | 0.08 ± < 0.01 |
| **Half-Life** | 12.83 ± 2.68 | 7.33 ± 0.43 | 10.72 ± 1.28 | 8.31 ± 0.55 |
| **Tau** | 18.51 ± 3.87 | 10.58 ± 0.62 | 15.47 ± 1.85 | 11.99 ± 0.80 |
| **Span** | 101.37 ± 55.61 | 332.23 ± 45.16 | 144.04 ± 41.56 | 251.10 ± 51.19 |

**Supplementary Table 7:** Average collagen fibre parameters extracted from SHG images of bone and osteosarcoma biopsies of increasing cancer stage. Data represents the mean of three independent samples ± standard deviation.

| Collagen Fibre Parameter | Tissue Type | | | | |
|---|---|---|---|---|---|
| | Bone | OS IB | OS IIB | OS III | OS IVB |
| Average Density (fibres / mm²) | 4470 ± 358.60 | 4284 ± 257.60 | 3055 ± 1573 | 4903 ± 137.10 | 4888 ± 248.20 |
| Average Straightness (p.d.u) | 0.91 ± < 0.01 | 0.91 ± < 0.01 | 0.91 ± < 0.01 | 0.89 ± < 0.01 | 0.89 ± < 0.01 |
| Average Width (μm) | 2.16 ± 0.03 | 1.82 ± 0.06 | 1.97 ± 0.36 | 1.82 ± 0.08 | 1.76 ± 0.13 |
| Average Length (μm) | 27.88 ± 2.18 | 23.94 ± 1.47 | 24.31 ± 0.93 | 23.16 ± 0.44 | 23.34 ± 0.85 |
| Av. Top 50th Percentile (μm) | 37.37 ± 4.13 | 26.83 ± 2.59 | 30.68 ± 1.66 | 28.57 ± 0.79 | 28.92 ± 1.47 |
| Av. Lowest 50th Percentile (μm) | 18.40 ± 0.24 | 17.91 ± 0.36 | 17.93 ± 0.22 | 17.75 ± 0.08 | 17.78 ± 0.26 |
